# Analytical validation and performance characteristics of a 48-gene next-generation sequencing panel for detecting potentially actionable genomic alterations in myeloid neoplasms

**DOI:** 10.1101/2020.11.30.403634

**Authors:** Sun Hee Rosenthal, Anna Gerasimova, Charles Ma, Hai-Rong Li, Andrew Grupe, Hansook Chong, Allan Acab, Alla Smolgovsky, Renius Owen, Christopher Elzinga, Rebecca Chen, Daniel Sugganth, Tracey Freitas, Jennifer Graham, Kristen Champion, Anindya Bhattacharya, Frederick Racke, Felicitas Lacbawan

**Affiliations:** Department of Advanced Diagnostics, Quest Diagnostics, San Juan Capistrano, CA; Department of Molecular Oncology, med fusion, Lewisville, TX

## Abstract

Identification of genomic mutations by molecular testing plays an important role in diagnosis, prognosis, and treatment of myeloid neoplasms. Next-generation sequencing (NGS) is an efficient method for simultaneous detection of clinically significant genomic mutations with high sensitivity. However, due to lack of standard NGS protocols, the application of NGS for hematologic malignancies into clinical settings remains limited. We report development and validation of a 48-gene NGS panel for molecular profiling of myeloid neoplasms including acute myeloid leukemia (AML), myelodysplastic syndrome (MDS), and myeloproliferative neoplasms (MPN). Target regions were captured by hybridization with complementary biotinylated DNA baits, and NGS was performed on an Illumina NextSeq500 instrument. A bioinformatics pipeline that was developed in-house was used to detect single nucleotide variations (SNVs), insertions/deletions (indels), and *FLT3* internal tandem duplications (*FLT3*-ITD). An analytical validation study was performed on 184 unique specimens for variants with allele frequencies ≥5%. Variants identified by the 48-gene panel were compared to those identified by a 35-gene hematologic neoplasms panel using an additional 137 unique specimens. The developed assay was applied to a large cohort (n=2,053) of patients with suspected myeloid neoplasms. Analytical validation yielded 99.6% sensitivity (95% CI: 98.9-99.9%) and 100% specificity (95% CI: 100%). Concordance of variants detected by the 2 tested panels was 100%. Among patients with suspected myeloid neoplasms (n=2,053), 54.5% patients harbored at least one clinically significant mutation: 77% in AML patients, 48% in MDS, and 45% in MPN. Together, these findings demonstrate that the assay can identify mutations associated with diagnosis, prognosis, and treatment options of myeloid neoplasms.

## Introduction

Myeloid neoplasms are a heterogeneous group of malignancies of the hematopoietic stem/progenitor cells. Substantial clinical and genomic overlap exists among different subclasses of myeloid neoplasms that are currently classified by the World Health Organization (WHO): acute myeloid leukemia (AML), myelodysplastic syndrome (MDS), myeloproliferative neoplasms (MPN), and overlap myelodysplastic/myeloproliferative neoplasms (MDS/MPN) [1]. Coupled with clinical, morphologic, and immunophenotypic abnormalities, identification of genetic alterations by molecular testing has an important role in the classification, risk stratification, and management of myeloid neoplasms [2]. For example, mutations in *NPM1, CEBPA,* and *RUNX1* can identify specific subclass of AML, and mutations in *FLT3, IDH1, IDH2,* and *JAK2* can guide targeted therapies for AML [3]. For patients with MDS, identification of mutations in genes including *TET2, SF3B1, ASXL1* and *TP53* is particularly useful to help establish clonal hematopoiesis to make a definitive diagnosis of MDS [4]. In addition, for MPN, certain mutations such as *JAK2* V617F or exon 12 mutation satisfy diagnostic criteria to help establish a diagnosis of MPN [5]; mutations in *ASXL1, SRSF2, EZH2, IDH1,* and *IDH2* categorize patients to high molecular risk in Primary Myelofibrosis (PMF) [6]; and mutations in *IDH2, U2AF1, EZH2, TP53, SH2B3*, and *SF3B1* indicate adverse prognostic value in Essential Thrombocythemia (ET) and Polycythemia Vera (PV) [7, 8].

Historically, single gene testing using Sanger sequencing or real-time PCR have been used to identify genetic alterations in myeloid neoplasms [9]. Although these methods are readily adaptive and widely used in clinical laboratories, their major limitation is that very few selected mutations can be detected at a time. In recent years, whole genome sequencing (WGS) and whole exome sequencing (WES) studies have been used to identify oncogenic mutations in hundreds of genes, revealing a wide genetic heterogeneity in myeloid neoplasms [9]. Indeed, more molecular genetic markers have been added in the most recent revision of the WHO classification [1]. Since the number of mutations that can guide diagnosis, prognosis, and treatment options are increasing, using single-gene testing for myeloid malignancies is becoming impractical. Although WGS and WES are standard in research settings, targeted next generation sequencing (NGS) panel assays that are composed of genes associated with a disease and recurrently mutated are more commonly used in clinical settings. Compared to WGS or WES, targeted NGS assays are highly sensitive for detecting low-frequency variants and can identify a number of mutations that are critical in diagnosis and risk stratification in a relatively short time [10].

Although an increasing number of hematologic laboratories are in the process of integrating NGS procedures into the diagnostic algorithms of myeloid neoplasms, the application of NGS in clinical settings has certain challenges [11]. For example, many artifacts are known to arise during NGS library preparation, sequencing, and data analysis (eg, read mapping, variant calling) and these may cause challenges in discriminating true genetic alterations from artifacts caused by PCR, sequencing, and post-sequencing steps. In addition, technical difficulties hinder the ability to capture targets with high GC content, such as *CEBPA,* which is associated with poor prognosis of AML, or repetitive genomic regions, such as *FLT3*-ITD, which is associated with poor prognosis of AML in the absence of *MPN1* mutation [10, 12, 13–15]. The lack of uniform practice standards for quality assessment of NGS data also challenges implementation of NGS in clinics [11]. As such, it is critical to carefully validate each clinical NGS assay for its defined performance requirements for the intended use.

In this study, we report development and validation of a 48-gene NGS panel for the detection of alterations that have a putative role in diagnosis, prognosis, or therapy of myeloid neoplasms. We also report our experience with the first 2,053 clinical specimens of suspected myeloid neoplasms submitted for the 48-gene panel and estimate the frequency of actionable alterations in a clinical laboratory.

## Materials and Methods

### Gene selection

A total of 48 genes (**S1 Table**) frequently mutated and/or associated with known diagnostic, prognostic, or therapeutic utilities for myeloid neoplasms at the time of test development were selected: 42 genes for AML; 36 genes for MDS; and 26 genes for MPN. Of the 48 genes, 47 were analyzed by NGS: the entire coding regions were covered for 23 genes (*ATM, BCOR, BCORL1, BRAF, CDKN2B, CEBPA, CREBP[ATF2], CUX1, DDX41, ETNK1, ETV6, GATA2, HRAS, IKZF1, KDM6A, NF1, PHF6, PTEN, STAG2, STAT3, STK11, TP53,* and *ZRSR2*) and targeted exons were covered for 24 genes *(ASXL1, CALR, CBL, CSF3R, DNMT3A, EZH2, FLT3 GATA1, IDH1, IDH2, JAK2, KIT, KRAS, MPL, NPM1, NRAS, PTPN11, RUNX1, SETBP1, SF3B1, SRSF2, TET2, U2AF1,* and *WT1). KMT2A* (formerly known as *MLL*) partial tandem duplication (PTD) analysis was supplemented by a long-range PCR method developed in our laboratory using a previously reported primer set, a forward primer on exon 8 and a reverse primer on exon 2 [16], to detect the 3 most frequent forms of *MLL*-PTD (NM_005933.1: exons 2~8, 2~9, and 2~10) [17].

### Validation specimens

A total of 184 unique specimens (**S2 Table**) were included: 96 whole blood, 20 bone marrow aspirate, 20 cell pellet, 17 FFPE, and 31 extracted DNA. The 184 specimens included 32 commercial controls, 25 from healthy individuals, and 127 clinical specimens previously tested by alternative analytical methods for indications of myeloid neoplasms (n=105), unknown history (n=18), and adjacent normal tissue from FFPE section (n=4). All patient specimens were de-identified before use. Alternative analytical methods included Sanger sequencing, fragment analysis for *FLT3*-ITD, and 4 different CLIA-validated NGS assays developed in our laboratory (see Alternative analytical methods, below). The 32 commercial controls included 6 multiplex mutation controls (5% Tier, Horizon discovery, Lafayette, CO), 19 controls with known mutation(s) within the 48 genes in the panel (Coriell Institute, Camden, NJ), and 7 well-characterized Genome In a Bottle (GIAB) reference specimens (Coriell Institute, Camden, NJ). For the 7 GIAB specimens, high-quality public sequence data were downloaded from http://ftp-trace.ncbi.nlm.nih.gov/giab/ftp/release/ for comparison.

### Alternative analytical methods

Sanger sequencing on *CEBPA* or *JAK2* were performed using gene-specific PCR primers for target amplification and incorporation of BigDye Terminator (Applied biosystems, Foster City, CA, USA). T7 promoter and terminator sequences fused to gene specific primers were used for sequencing on an ABI 3730 fragment analyzer (Applied biosystems, Foster City, CA, USA).

*FLT3*-ITD fragment analysis was performed by Laboratory for Personalized Molecular Medicine (LabPMM, San Diego, CA) or in-house using a modified protocol based on one reported previously [18]. In brief, *FLT3*-ITD was PCR-amplified with a fluorescence-labeled forward primer, and a non-labeled reverse primer. The PCR products were analyzed using an ABI3730 genetic analyzer (Applied Biosystems, Foster City, CA, USA), and the amplicons with a size greater than that of wild type (324-326 bp) were interpreted as positive for the *FLT3*-ITD.

Four separate CLIA-validated NGS panel assays were used for the accuracy study to cover the variants detected by the 48-gene panel. The first, LeukoVantage v1.0, is a 30-gene NGS test for somatic mutations in myeloid neoplasms using the Truseq Amplicon Cancer Panel kit (Illumina, San Diego, CA). It contains amplicons for hot-spot locations of the following genes: *ASXL1, CALR, CBL, CEBPA, CSF3R, DDX41, DMNT3A, EZH2, FLT3 GATA1, IDH1, IDH2, JAK2, KDM6A, KIT, KRAS, MLL, MPL, NPM1, NRAS, PTPN11, RUNX1, SETBP1, SF3B1, SRSF2, TET2, TP53, U2AF1, WT1*, and *ZRSR2* [19]. The second, MyVantage, is a germline mutation panel of 34 cancer predisposition genes, including *ATM* and *PTEN,* and uses RNA bait capture method [20]. The third, Watson Genomics, is a 50-gene NGS test for solid tumor mutation using RNA bait capture-based NGS, and includes *BRAF, CDKN2B, FLT3, HRAS, IDH1, JAK2, KIT, KRAS, NRAS, PTEN, PTPN11*, and *TP53*. The fourth, MPN diagnostic cascading reflex test, includes hotspots in *MPN, CALR, MPL*, and *CSF3R* genes and employs multiplex PCR method and sequencing on Ion Torrent S5 XL (Thermo Fisher Scientific, Markham, ON).

### DNA isolation, library preparation, and NGS

Genomic DNA was isolated from whole blood, bone marrow, or cell pellet using Qiagen EZ1 kit or DSP DNA mini kit (Qiagen, Mississauga, ON). FFPE DNA was extracted using QIAAMP DNA FFPE tissue kit (Qiagen, Mississauga, ON), or Arcturus PicoPure kit (Thermo Fisher Scientific, Markham, ON). Isolated genomic DNA was mechanically sheared to an average size of 250 bases using a Covaris instrument LE220 (Covaris Inc., Woburn, MA). The fragmented DNA was enzymatically repaired and end-modified with adenosine (NEBNext^®^ Ultra™ II End Repair/dA-Tailing Module, NEB, Ipswich, MA) and ligated (NEBNext® Ultra ™ II Ligation Module, NEB, Ipswich, MA) with barcoded adapters (Integrated DNA Technologies, Coral, IL). The ligated products were size-selected (AMPure Beads, Agencourt, Beverley, MA) and amplified (GeneRead DNA I Amp Kit, Qiagen, Mississauga, ON); then the regions of interest were captured using biotinylated DNA baits (Integrated DNA Technologies, Coral, IL). The hybridized DNA fragments were enriched with streptavidin-attached magnet beads (Dynabeads M-270, Thermo Fisher Scientific, Markham, ON) and washed under increasing stringency to remove non-targeted DNA sequences (xGen® Lockdown® Reagents, Integrated DNA Technologies, Coral, IL). A second amplification was performed (KAPA HiFi HotStart ReadyMix, Kapabiosystems, Wilmington, MA), followed by bead purification (AMPure Beads, Agencourt, Beverley, MA) to remove all unused primers and nucleotides. The prepared sequencing library was then quantified (Qubit dsDNA HS Assay Kit, Thermo Fisher Scientific, Markham, ON) and sequenced on an Illumina NextSeq 500 sequencer, 2 x 150 cycles (NextSeq 500/550 Mid Output v2 kit, Illumina, San Diego, CA).

### Bioinformatics processing

De-multiplexing and conversion of NextSeq500 BCL files were done by using Illumina’s bcl2fastq software utility. The raw sequence reads in FASTQ files were then aligned to the Genome Reference Consortium human genome build 37(GRCh37) by using BWA alignment package. Reads were sorted and indexed using SAMtools with subsequent reads duplication removal by Picard Tools. Local realignment and base quality score recalibration were performed using the Genome Analysis Toolkit (GATK). Mapped reads were further filtered by mapping score ≥ 30 (≥ 99.9% accuracy) and base quality score ≥ 20 (≥ 99% accuracy) before downstream analysis. This assay covered 98,809 bp from a total of 999 target regions, approximately100 bp for each target, across the 48 genes in the panel. Average and minimum depth of coverage for every region of interest (ROI) and depth of each targeted positions were computed using SAMtools Pysam. SNVs and short indels were called by MuTect2 and LoFreq. *CALR* indels and *FLT3*-ITD were called by PINDEL. Criteria used for specimen and variant quality control are provided in **S3 Table**.

### Variant call comparison

An additional 137 unique specimens that were submitted for the 48-gene panel or a 35-gene panel, which was independently developed and analytically validated for hematologic neoplasms (HemeSEQ, med fusion, Lewisville, TX), were subsequently sequenced by the other panel. The 35-gene panel uses Illumina TruSeq custom-amplicon library preparation chemistry, which is sequenced on an Illumina MiSeq. A total of 33,812 bp in 26 genes overlapped between the 2 assays; genes included *ASXL1, CALR, CBL, CSF3R, DNMT3A, ETV6, EZH2, FLT3, IDH1, IDH2, JAK2, KIT, KRAS, MPL, NPM1, NRAS, PHF6, RUNX1, PTEN, SETBP1, SF3B1, SRSF2, TET2, TP53, U2AF1,* and *WT1.* Each of the 137 specimens (22 whole blood and 115 bone marrow) were selected because they harbored at least one pathogenic mutation within the overlapping genes.

### Clinical specimens

A total of 2,053 consecutive patient specimens submitted for the 48-gene NGS panel were included in this study. Patient results were de-identified before analysis. Based on the clinical information submitted, indications for testing included AML (23.9%, n=490), MDS (49.6%, n=1,018), or MPN (26.5%, n=545). Patient characteristics are presented in **Table 1**. This retrospective study was exempt from Institutional Review Board oversight, as determined by the Western Institutional Review Board.

**Table 1.** Patient characteristics

|  | AML | MDS | MPN | Total |
| --- | --- | --- | --- | --- |
| Number of patients | 490 | 1,018 | 545 | 2,053 |
| % of total | 23.9 | 49.6 | 26.5 | 100.0 |
| Male, % | 61.6 | 61.5 | 57.8 | 60.5 |
| Female, % | 38.4 | 38.5 | 42.2 | 39.5 |
| Median age (Min, Max) | 58 (2, 89) | 60 (2, 89) | 55.5 (14, 69) | 58 (2, 89) |

### Variant classification

For the 2,053 clinical cases, clinical interpretation of variants was performed in conjunction with a third-party annotation group (N-of-One, Concord, MA). Detected variants were classified in the following categories based on the 2017 guideline recommendations by AMP/ASCO/CAP [21]: Tier I, strong clinical significance; Tier II, potential clinical significance; Tier III, uncertain clinical significance; and Tier IV, benign or likely benign. A pathologist or a licensed clinical laboratory director reviewed the results and the clinical annotation, edited information as deemed necessary for specific cases and indications, and signed out the results for reporting. For this study, variants were considered clinically significant if they provided prognostic or diagnostic information for the disease or were clinically actionable. Variants were considered clinically actionable when a targeted therapy or an experimental drug was available for the disease or other disease.

## Results

### Assay performance

The analytical validation study was performed on 184 unique specimens for a total of 427 trials in 20 consecutive sequencing runs. On average, each specimen generated 11.8 million reads; 100% (SD=8.8%) of reads mapped to the reference sequence (hg19); 65% (SD=8.8%) of reads were on-target; and average coverage depth across target regions was 1,767 (SD=875) (**S4 Table**). Average and median coverage of each target region across all validation specimens are plotted in **Fig 1A**. Of the 184 specimens, 181 (98.3%) specimens exceeded the coverage requirement in 422 trials (98.8%) and 3 specimens (1 whole blood, 1 FFPE, and 1 extracted DNA) failed in 5 trials. This result demonstrated that the developed workflow is robust for routine clinical testing and compatible with different specimen types.

**Fig 1.** Target coverage depth. (A) Mean and median coverage of each target region across all validation trials (n=427). Target regions are sorted by chromosomal location. Targets on X chromosome are marked with a dotted line. (B) NA12878 coverage across all target regions from 17 independent setups. Target regions are sorted by % GC (orange dot). Standard deviation of coverage is shown (black bar). (C) Percentage of specimens that passed a base quality Q20 coverage of 100X, 250X, or 500X.

We next assessed coverage at each target region as a function of target region % GC using a control specimen NA12878, which was tested 17 times. Throughout the whole % GC spectrum (range: 12%-87%), we observed narrow coverage distribution, even in the extreme end of % GC (**Fig 1B**). In addition, we reviewed specimen pass rates of each target for Q20 coverage of 500X, 250X, or 100X (**Fig 1C**). A limited number of target regions did not achieve 500X coverage in >10% of validation specimens. Those regions included *STAG2* exon 11, 16, and 21, all of which were situated near polyA tracts and represented 0.36% (354/98,809bp) of our target capture region. All 3 target regions achieved 100X coverage in 100% of validation sets and 250X coverage in >96% of validation sets.

### Precision

To test inter-assay precision, 22 specimens that harbored at least one pathogenic mutation by alternative methods and a negative control (NA12878) were analyzed 3 times in 3 independently prepared sequencing runs. A total of 141 variants (98 SNVs, 39 indels, and 4 *FLT3*-ITD) were detected and were 100% concordant among 3 repeated runs (**S5A Table**). As expected, no reportable variant was detected from the negative control (NA12878) after applying variant filtering rules (**S3 Table**). Intra-assay precision was tested on 9 specimens that were tested in triplicate within a run. A total of 40 variants (30 SNVs, 8 indels, and 2 *FLT3*-ITD) were detected and all were concordant among the triplicates (**S5B Table**). In addition, detected variant frequency among replicates was reproducible within coefficient of variation <0.2 in 98.3% and <0.25 in 100% of variants (n=181) tested in the precision studies (**Figs 2A and 2B**). In summary, both intra- and inter-assay variant call concordance was 100%.

**Fig 2.** (A) Inter-assay precision of 141 variants from 22 specimens repeated 3 times. The mean of the detected variant frequencies (orange dot) with standard deviation (closed vertical bar) and coefficients of variations (blue dot) are shown. (B) Intra-assay precision of 40 variants from 9 specimens replicated 3 times. The mean of the detected variant frequencies (orange dot) with standard deviation (closed vertical bar) and coefficients of variations (blue dot) are shown.

### Accuracy

A total of 140 unique specimens harboring variant(s) identified by alternative analytical methods, such as Sanger sequencing, fragment analysis, and other CLIA validated NGS assays, were analyzed for a total of 237 trials (**S6 Table**). The identified 165 unique variants by alternative methods consisted of 97 SNVs, 49 indels (1-33 bp insertions, 1-52 bp deletions), and 19 *FLT3*-ITDs (18-117 bp). From the 237 trials, collectively, 405 variants of various frequencies were analyzed including 229 SNVs, 117 indels, and 59 *FLT3*-ITDs in 29 genes (**Figs 3A, 3B**). All expected SNV and indel variants were accurately detected. If variant frequency was provided by an alternative method (n=106), frequency of detected variant was highly concordant with expected (R^2^=0.966, **Fig 3C**). In addition, all expected *FLT3*-ITD variants from 59 trials of 27 unique specimens were detected by this assay (**Table 2**). In 18.5% (5/27) of *FLT3*-ITD positive specimens, additional ITD sizes that were not reported by a fragment analysis method were detected; this result may indicate higher sensitivity of NGS assay for *FLT3*-ITD detection.

**Table 2.** *FLT3*-ITD used for accuracy study

| Sample | *FLT3*-ITD length (bp) | Number trial(s) | Detected | Additional *FLT3*-ITD detected (bp) |
| --- | --- | --- | --- | --- |
| BM08 | 36 | 3 | yes | - |
| BM15 | 57 | 3 | yes | - |
| BM17 | 57 | 3 | yes | - |
| BM19 | 60 | 3 | yes | - |
| CP04 | 45 | 1 | yes | - |
| CP17 | 24 | 1 | yes | - |
| CP20 | 21, 27, 36, 117 | 1 | yes | 48 |
| WB09 | 30 | 11 | yes | 33, 81 |
| WB32 | 21 | 1 | yes | - |
| WB45 | 96 | 6 | yes | - |
| WB46 | 72 | 3 | yes | - |
| WB47 | 51, 114 | 4 | yes | - |
| WB48 | 63 | 1 | yes | - |
| WB50 | 18, 63, 90 | 4 | yes | - |
| WB51 | 33, 57 | 1 | yes | - |
| WB52 | 21, 51, 90 | 1 | yes | - |
| WB53 | 18 | 1 | yes | - |
| WB54 | 36 | 1 | yes | 24 |
| WB56 | 24 | 1 | yes | - |
| WB57 | 48 | 1 | yes | 33 |
| WB58 | 30 | 1 | yes | - |
| WB59 | 36 | 1 | yes | - |
| WB60 | 51, 54 | 1 | yes | - |
| WB61 | 24 | 2 | yes | - |
| WB62 | 66 | 1 | yes | - |
| WB63 | 24 | 1 | yes | - |
| WB64 | 84 | 1 | yes | 27 |

BM, bone marrow; CP, cell pellet; WB, whole blood

**Fig 3.** (A) Number and type of variant used for accuracy study per gene. (B) Variant frequency distribution used for accuracy study. Variant frequency not provided by alternative method is categorized as undetermined. (C) Variant frequency concordance. Compared only if variant frequency is known by alternative method.

We extended the accuracy study to 7 well-characterized GIAB reference specimens by comparing variant calls made by our pipeline with publicly available data. The variant concordance study was limited to those regions where high-quality public sequence data were available, approximately 62% to 86% of our target regions for each GIAB reference specimen (**Table 3**). Population SNVs that would normally be excluded from reporting were also included for this study. Across the 7 reference specimens, a total of 464,162 bp were analyzed. All 142 expected variants were correctly detected (**Table 3**), and no false positives were called, yielding 100% sensitivity (95% CI: 97.3-100%) and 100% specificity (95% CI: 100%). Review of the detected variant frequency at the expected homozygous and heterozygous positions revealed the mean observed variant frequency to be 99.9% (N=70, SD=0.1) and 49.6% (N=72, SD=2.6), respectively, demonstrating good agreement with the expected values.

**Table 3.** Variant call concordance study using Genome in a Bottle reference specimens

| **Specimen** | **Overlapping regions:**<br>**High confidence sequence from**<br>**GIAB** | **Number of**<br>**variants** | **% Concordance** |
| --- | --- | --- | --- |
| NA12878 | 86,423 bp | 23 | 100 |
| NA24143 | 62,850 bp | 16 | 100 |
| NA24149 | 62,924 bp | 20 | 100 |
| NA24385 | 62,815 bp | 22 | 100 |
| NA24631 | 63,047 bp | 21 | 100 |
| NA24694 | 63,070 bp | 19 | 100 |
| NA24695 | 63,033 bp | 21 | 100 |
| Total | 464,162 bp | 142 | 100 |

### Analytical sensitivity

In order to evaluate the analytic sensitivity of the assay, 5 multiplex mutation control specimens (5% Tier, Horizon discovery) were tested for a total of 16 trials. Collectively, 124 expected variants (104 SNVs and 20 indels) with variant allele frequency (VAF) ranging from 3.8% to 25% were identified (**S7A Table**). All expected variants were detected when expected VAF was either at 5% (71/71) or >5% (35/35). However, 89% (16/18) of expected variants were detected when expected VAF was <5%.

In addition, DNA from each of 5 well-characterized GIAB reference specimens was mixed with NA12878 DNA in a proportion of 10% test specimen and 90% NA12878. From heterozygous or homozygous variants unique to test specimens, a total of 64 variants (63 SNVs and 1 insertion) were created with VAF 10% or 5% (**S7B Table**). A total of 119 variants were tested from the 5 mixed specimens of which 4 specimens were repeated twice. All (50/50) variants were detected at 10% VAF and 95.6% (66/69) variants were detected at 5% VAF.

Similarly, 9 specimens harboring variants (2 SNVs and 10 indels) identified by alternative analytical methods were serially diluted with a GIAB control (NA12878) up to 64-fold, yielding an expected VAF well below 5% (**S7C Table**). All expected variants were detected when expected VAF was >5% (n=45). However, when expected VAF was below 5%, only 41% (2/8 at <3.0%, 5/9 at 3.1-3.4%) of expected variants were detected.

In addition, 3 *FLT3*-ITD positive specimens were serially diluted up to 16-fold with NA12878. We considered *FLT3*-ITD positive when at least one ITD size is called. All dilution series yielded correct ITD calls (**S7D Table**). As the orthogonal method does not provide absolute *FLT3*-ITD frequency, in order to estimate sensitivity, we simulated expected variant frequency based on the relative fraction of different ITD sizes in a specimen, WB52. WB52 contained 3 different ITD sizes (21b, 51b, and 90b) with relative ITD fraction of 1.4, 75.7, and 23.0%. The major ITD size (51b) was detected from all dilution series including 16-fold dilution with simulated VAF of 4.7% (16-fold dilution of 75.7%). This result demonstrated that *FLT3*-ITD detection sensitivity to be at least 5%.

A summary of analytical sensitivity study results is provided in **Table 4**. Inspection of the discordant variants from this study (12 at ≤3.1% VAF and 3 at 5% VAF) showed that those variants had been detected by a variant caller, but the calls were filtered out because of low frequency (<3%) or low variant count (<25) (**S8 Table**).

**Table 4.** Summary of analytical sensitivity study result

| Study | Expected VAF |  |  | Total |
| --- | --- | --- | --- | --- |
|  | <5% | 5% | >5% |  |
| Multiplex mutation control | 89 (16/18) | 100 (71/71) | 100 (35/35) | 98 (122/124) |
| GIAB dilution | N/A | 96 (66/69) | 100 (50/50) | 97 (116/119) |
| Positive specimen dilution | 41 (7/17) | N/A | 100 (58/58) | 87 (65/75) |
| Total | 66 (23/35) | 98 (137/140) | 100 (143/143) | 95 (303/318) |

Performance is provided as % detected (no. detected/no. expected). N/A, not applicable.

Collectively, in this validation, a total of 154 unique specimens were analyzed for 830 known variants (588 SNVs, 167 indels, and 75 *FLT3*-ITDs) of VAF ≥5%, including results described in the accuracy section. Almost all variants (99.6%, 827/830) of VAF ≥5% variants were correctly detected, resulting in analytical sensitivity of 99.6% (95% CI: 98.9-99.9%). Analytical sensitivity for various VAFs for each variant type is summarized in **Fig 4** and **S9 Table**. In conclusion, the variant detection limit for this assay is 5% for SNV, indels (including *CALR* 52 bp deletion), and *FLT3*-ITD.

**Fig 4.** Analytical sensitivity of the assay for various VAFs for each variant type. Analytical sensitivity percentage represents the proportion of detected variants out of all variants tested at a given VAF.

### Variant call comparison study

To increase per-sample data comparison efficiency, we compared variants of an additional 137 unique specimens identified by the validated 48-gene NGS panel for myeloid neoplasms to those detected by an independently developed and analytically validated 35-gene NGS panel for hematologic neoplasms (HemeSEQ, med fusion, Lewisville, TX). The 137 specimens were selected because they harbored at least one pathogenic mutation within overlapping genes (n=26) between the 2 assays. From the 137 specimens (141 trials), a total of 1,094 variants (278 unique) of various frequencies were concordantly detected by the 2 assays: 1,007 (219 unique) SNVs and 87 (59 unique) indels (**Fig 5A**). In this study, benign variants (eg, common population SNVs) were also included. In addition, agreement in VAF between the 2 assays was good (R^2^=0.986) (**Fig 5B)**. Collectively, a total of 4.8 million individual base calls (33,812 bp X 141 specimens) were compared, including 1,094 variants and resulted in 100% sensitivity (95% CI: 99.7-100%) and 100% specificity (95% CI: 99.9-100%).

**Fig 5.** Variant call concordance study between the 48-gene panel for myeloid neoplasms and 35-gene panel for hematologic neoplasms. (A) Number and type of variants compared per gene. (B) Scatter plot of VAFs (n=1,094) across 141 trials detected by the 48-gene panel (x-axis) and 35-gene panel (y-axis).

### Long-term test reproducibility

To evaluate long-term reproducibility, we examined variant calls for a multiplex mutation control (HD728) that was repeatedly tested, 119 times over 10 months, as a mutation positive control. The mutation control contained 7 alterations of ≥5% VAF (6 SNVs and 1 deletion, **Table S7A**). All alterations were successfully detected in all repeat tests, including 5 alterations occurring at 5% VAF. Detected VAF remained stable over the extended time period (**Fig 6**) with coefficient of variation less than 0.1x (range: 0.04-0.1x).

**Fig 6.** Detected variant frequency of a mutation-positive control (HD728) from 119 repeat tests over 10 months. Expected variant frequency is indicated along with variant name.

### Detection of clinically actionable alternations

Retrospective analysis was performed on the mutation profiling of 2,053 consecutive, deidentified unique patient specimens submitted for the 48-gene NGS panel. In total, 98.3% (2,018/2,053) of specimens were successfully tested upon initial testing and the remaining 35 specimens passed QC criteria upon repeat test. Based on the clinical information submitted, indications for testing included 490 AML (23.9%), 1,018 MDS (49.6%) and 545 MPN (26.5%). At least one pathogenic mutation was detected in 55.6% (1,142/2,053) of patient specimens. AML patients had the highest positive rate (81.2%, 398/490), followed by MDS (49.0%, 499/1,018) and MPN (45.0%, 245/545). A median of 2 mutations (range 1-12) were detected per positive patient: 3 (1-12) in AML, 2 (1-9) in MDS, and 2 (1-7) in MPN. Collectively 2,799 pathogenic mutations (Tier I or Tier II) were found in 44 genes (**Fig 7**). The most frequently mutated genes, which were found in at least 10% of patients for each indication, were *TET2, ASXL1, TP53, FLT3, NPM1, DNMT3A, IDH2, RUNX1* and *NRAS* in AML; *TET2, SF3B1,* and *ASXL1* in MDS; and *JAK2, TET2,* and *ASXL1* in MPN (**S1 Fig)**. The genomic alterations identified across 2,023 specimens are depicted in **Fig 8**.

**Fig 7.** Pathogenic mutations (Tier I or Tier II) detected in 44 genes from the total cohort of 2,053 patients by the 48-gene NGS panel for myeloid neoplasms.

**Fig 8.** Identified Tier I and Tier II mutations in 44 genes from the total cohort of 2,053 patients detected by the 48-gene NGS panel for myeloid neoplasms. Each column in the x=axis represents a patient. Only patients with at least 1 mutation (n=1,142) are shown. The percent of patients with Tier I and Tier II mutations in the indicated gene is presented.

Based on functionally related categories (**Fig 8**), genes involved in epigenetics (*ASXL1, BCOR, BCROL1, DNMT3A, EZH2, IDH1, IDH2, KDM6A,* and *TET2*) were the most frequently mutated group, detected in 43% (n=875/2,053) of patients. Genes involved in signal transduction (*BRAF, CALR, CBL, CSF3R, FLT3, JAK2, KIT, KRAS, MPL, NF1, NRAS, PTPN11*, and *STAT3*), RNA splicing (*SF3B1, SRSF2, U2AF1*, and *ZRSR2*), and transcription factor (*CEBPA, ETV6, GATA1, GATA2, IKZF1, KMT2A, PHF6, RUNX1, SETBP1,* and *WT1)* accounted for 27% (n=574/2,053), 22% (n=444/2,053), and 13% (n=272/2,053) of patients, respectively. These findings are consistent with the mutational frequency of gene groups in myeloid neoplasms as reported in literature [22–26].

In 41.7% (856/2,053) of patients, at least 1 actionable mutation was identified (**Fig 9**): 27.5% (n=565) of patients harbored mutations for which a targeted therapy is available either in the disease (n=126) or another disease (n=439); and 40.1% (n=823) patients contained mutations for which an experimental drug is available. For this study, variants were considered clinically actionable when a targeted therapy or an experimental drug was available for the disease or other disease. In addition, 34.6% of patients (n=711) had mutations with prognostic significance, and 36.4% (n=748) of patients had mutations with diagnostic significance. In total, the assay identified clinically significant mutations in 51.7% (1,062/2,053) of patients. Mutations were considered clinically significant if they provided prognostic or diagnostic information for the disease or were clinically actionable.

### Discussion

Molecular profiling can help diagnose, classify, and guide treatment of myeloid neoplasms [27–28]. In this study, we reported development and validation of a 48-gene NGS panel for molecular profiling of myeloid neoplasms. The assay demonstrated good inter- and intra-assay precision for SNVs, indels including *CALR* 52 bp deletion, and *FLT3*-ITDs (**Fig 2**, **S5A and S5B Tables**). In addition, the assay detected 827 of 830 variants with VAF ≥5% reported by alternative analytical methods for a sensitivity of 99.6% (95% CI: 98.9-99.9%) (**Fig 4**, **S8 Table**). False positive calls with low frequency in low complexity sequence regions are a relatively common phenomenon in NGS [11]. The possibility of detecting this type of event has been reduced by using bioinformatic filtering process established based on the results of a training set of 100 unique normal patient specimens (data not shown). The specificity of this assay appears to approach 100% as no false positive call was made in 7 GIAB accuracy studies across 0.4 million bp (**Table 3**) and in a cross-platform study using 137 unique specimens across 4.8 million bp (**Fig 5**).

In clinical settings, specimen types submitted for an assay may vary. Thus, an ideal assay must be able to deal with a wide range of specimen types appropriate for the assay. Overall, our assay produced robust results (98.8% pass rate) for all specimen types (**Table S2**), suggesting that DNA recovered from various specimen types can be successfully and accurately sequenced by this protocol. In NGS, the limitation of a target enrichment method often leads to low coverage in genes with high GC content, and this may cause sub-optimal assay accuracy. Supplementary single-gene assays are recommended to improve low coverage targets in some clinical assays [29]. In this study, we demonstrated a broad reportable range of the 48-gene panel even in extreme % GC spectrum (**Fig 1B**). For example, despite the very high GC content of *CEBPA* gene (up to 87% GC within a 100-bp window), hybridization enrichment coupled with careful bait design achieved an excellent result: median depth of 1,918X. During our development, direct comparison of capture technologies demonstrated superior performance of DNA bait over RNA bait for targets with high GC contents (**S2 Fig**). During our validation, we showed that VAF of SNVs and indels detected by our assay were highly concordant with ones observed by reference methods (**Fig 3C**, **Fig 5B**), indicating our assay is accurate in variant quantification. Long-term test reproducibility of an analytical method is critical in clinical settings following initial validation; consistent variant allele frequencies over a 10-month period in a positive control specimen highlighted the long-term reproducibility of our assay (**Fig 6**). *FLT3*-ITD is inherently difficult to detect using NGS approaches [15, 30]. In our validation studies, we also identified *FLT3*-ITDs of varying lengths (range: 18-117 bp) with 100% sensitivity and specificity (**Table 2**). In some trials, additional ITD size(s) were detected, and we reasoned that our assay is more sensitive than the method compared, fragment analysis.

In our study of 2,053 clinical patients, 55.6% had at least one pathogenic variant and 51.7% harbored clinically significant mutations with prognostic, diagnostic, or therapeutic relevance (**Fig 9**). The clinical utility of our assay is underscored by our ability to identify clinically significant variants in specific diseases. For example, from our cohort of 490 patients with indications of AML, *TET2* had the highest mutation rate (17% of patients) followed by *ASXL1, TP53, FLT3, NPM1, SRSF2, DNMT3A, IDH2, RUNX1* and *NRAS* (10~15%), frequencies similar to literature [26]. Among those frequently mutated genes, *ASXL1, RUNX1, TP53,* and *FLT3*-ITD mutations have been associated with poor prognosis, whereas *NPM1* mutations in the absence of *FLT3*-ITD have been associated with favorable outcomes in AML [31]. In the 2016 revision of the WHO classification, AML with an *NPM1* mutation is recognized as a subtype of AML; and AML with an *RUNX1* mutation has been added as a provisional category of AML [1]. In addition, FDA-approved targeted therapies for AML are available for *FLT3*-ITD and *IDH2,* as well as *IDH1* [32] which was mutated in 5% of patients in our study.

From our cohort of 1,018 patients with indications of MDS, *TET2, SF3B1,* and *ASXL1* were the most frequently mutated genes (in >10% of patients), followed by *SRSF2, TP53, DNMT3A,* and *RUNX1* (in >5% of patients), similar to Haferlach and colleague’s study on the mutational profiles of 944 patients with MDS [25]. Among those genes, *ASXL1* and *TP53* mutations have been associated with poor prognosis [33], whereas *SF3B1* mutation in MDS patients with ringed sideroblasts has been associated with favorable prognosis [34]. All of the frequently mutated genes have been associated with clonal hematopoiesis, and support the diagnosis of several different myeloid malignancies, including MDS, when found in combination with other diagnostic features [35, 36]. While there are currently no therapies directly targeting mutated genes in MDS, hematologic malignancies harboring *SF3B1* and *SRSF2* mutations have been reported to be sensitive to splicing factor 3B subunit 1 (SF3b155) inhibitors, which are in clinical development [37, 38].

Certain mutations in the MPN-associated genes satisfy subclassification of the disease. In our study of 545 patients with indications of MPN, *JAK2* was the most frequently mutated gene (23% of patients) followed by *TET2* (12%) and *ASXL1* (10%), frequencies similar to literature [39, 40]. *CALR, CSF3R,* and *MPL* were mutated in 4%, 3%, and 1% patients, respectively. Presence of a mutation in *JAK2, CALR,* and *MPL* is among the major criteria for the diagnosis of myelofibrosis (MF) or essential thrombocythemia (ET), while presence of *JAK2* V617 or exon 12 mutations is among the major criteria for the diagnosis of polycythemia vera (PV) [1, 41, 42]. Activating *CSF3R* mutations have been found in the majority of chronic neutrophilic leukemias (CNLs) [43]. FDA-approved therapies for *JAK2* mutations in PV and MF are available [44]. In addition, mutations in *CALR* exon 9, *CSF3R,* and *MPL* have been shown to be sensitive to Jak inhibitors [45–47].

For *BCR*-*ABL1*-negative MPN, common mutations of *JAK2, CALR,* and *MPL* genes are often examined as diagnostic targets using a cascade single-gene assay [42]. In our cohort of 545 patients with indications of MPN, 159 patients (29%) were positive for those 3 genes. Detection of an additional clinically significant mutation is more common in an NGS panel assay than single-gene tests owing to the multiplicity of genes tested. An additional 86 patients (16%) who were negative for those 3 genes were positive for pathogenic mutations in other genes in the panel. Of the 86 patients, 23 had mutations in genes that can aid diagnosis when found in combination with other diagnostic features *(CBL, NRAS,* and *PTPN11)* [1, 8, 36], 43 had mutations in genes with poor prognosis (*ASXL1, EZH2, IDH2, NRAS, SETBP1, SRSF2, TP53*, and *U2AF1*), and 29 had mutations in genes for which a therapy is available (*BRAF, CSF3R, IDH2, KRAS, NF1, NRAS, PTPN11,* and *STAG2).* In addition, of the 159 patients who were positive for those 3 genes, 84 patients had additional mutation(s) in 16 other genes. Among the 16 genes, 6 genes *(IDH1, IDH2, KRAS, NF1, NRAS,* and *STAG2)* are linked to available therapies in other diseases [27, 48]. Combinations of Jak inhibitors with other targeted therapies may be relevant for those patients who harbor additional mutations.

In contrast to single-gene assays, NGS allows assessment of co-occurring mutations that might have heterogeneity of response to targeted therapy and survival. According to the recent National Comprehensive Cancer Network guidelines, AML patients with *FLT3*-ITD and *NPM1* double mutation (AML *FLT3*-ITD*+/NPM1+*) are categorized as favorable and intermediate risk levels, depending on the allelic ratio of *FLT3*-ITD, for whom allogenic stem cell transplantation (allo-HSCT) is not obligated. Loghavi and colleagues reported that AML *FLT3-ITD+/NPM1+* patients with a *DNMT3A* mutation had shorter event-free survival compared to those in other mutation groups [49]. Similarly, recent studies have suggested that AML patients with concomitant *DNMT3A R882+/FLT3-ITD+/NPM1+* mutations had a very poor prognosis, and allo-HSCT could moderately improve their survival [50, 51]. Our study cohort included 21 AML *FLT3-ITD+/NPM1+* patients, 4 of whom had a *DNMT3A* R882 mutation. In addition, Ardestani and colleagues reported that *DNMT3A* R882 mutations alone do not affect the clinical outcomes of AML patients, but when accompanied by *FLT3*-ITD mutations, overall survival was reduced, even after allo-HSCT [52]. In our study, 5 of 26 *DNMT3A* R882-positive AML patients had *FLT3*-ITD. These results support the clinical utility of our assay for detecting mutations that can alter prognostic and therapeutic significance when they occur in combination.

## Conclusions

We have developed and validated a 48-gene NGS assay that can detect SNVs, indels, and *FLT3*-ITD with high sensitivity and specificity. The assay detects variants with clinical significance from a substantial proportion of patients tested. The developed assay may be used to guide more precise and targeted therapeutic strategies, possibly leading better treatment outcomes for patients with myeloid neoplasms.

## Acknowledgments

Authors thank Kevin Qu and Quoclinh Nguyen for technical assistance; Joseph Catanese and Feras Hantash for technical supervision; Andy Le, Trina Nguyen, and Suzette Utulo for clinical data generation; Lucas Xu for data query; Andrew Hellman and Melissa Smith for critical review of the manuscript.

## Supporting information

**S1 Table. Genes included in the 48-gene NGS panel.**

**S2 Table. Specimens used for validation studies.**

**S3 Table. Criteria used for specimen and variant quality control.**

**S4 Table. Summary of sequencing metrics for validation specimens (n=428,184 unique).**

**S5A Table. Inter-assay precision study variant concordance.**

**S6 Table. Specimens used for accuracy study.**

**S7A Table. Variant call concordance study using multiplex mutation control specimens.**

**S7B Table. GIAB reference specimen dilution study.**

**S7C Table. Positive specimen dilution study.**

**S7D Table. *FLT3-ITD* positive specimen dilution study.**

**S8 Table. Summary of discordant variant.**

**S9 Table. Analytical sensitivity for various % VAF for each variant type.**

**S1 Fig. % patient with mutation per gene for AML (A), MDS (B), and MPN (C).** Pathogenic (Tier I and Tier II) mutations were included.

**S2 Fig. Integrative Genomics Viewer (IGV) for *CEBPA* (A) and *CUX1* (B) comparing target capture performance between RNA bait (top panel) and DNA bait (bottom panel).** Even coverage distribution was achieved using DNA bait whereas low to no coverage was observed using RNA bait for high % GC targets. % GC in 100bp window is color-coded.

